# Effect of lossy compression of quality scores on variant calling

**DOI:** 10.1101/029843

**Authors:** Idoia Ochoa, Mikel Hernaez, Rachel Goldfeder, Tsachy Weissman, Euan Ashley

## Abstract

Recent advancements in sequencing technology have led to a drastic reduction in the cost of genome sequencing. This development has generated an unprecedented amount of genomic data that must be stored, processed, and communicated. To facilitate this effort, compression of genomic files has been proposed. Specifically, lossy compression of quality scores is emerging as a natural candidate for reducing the growing costs of storage. A main goal of performing DNA sequencing in population studies and clinical settings is to identify genetic variation. Though the field agrees that smaller files are advantageous, the cost of lossy compression, in terms of variant discovery, is unclear.

Bioinformatic algorithms to identify SNPs and INDELs from next-generation DNA sequencing data use base quality score information; here, we evaluate the effect of lossy compression of quality scores on SNP and INDEL detection. We analyze several lossy compressors introduced recently in the literature. Specifically, we investigate how the output of the variant caller when using the original data (uncompressed) differs from that obtained when quality scores are replaced by those generated by a lossy compressor. Using gold standard genomic datasets such as the GIAB (Genome In A Bottle) consensus sequence for NA12878 and simulated data, we are able to analyze how accurate the output of the variant calling is, both for the original data and that previously lossily compressed. We show that lossy compression can significantly alleviate the storage while maintaining variant calling performance comparable to that with the uncompressed data. Further, in some cases lossy compression can lead to variant calling performance which is superior to that using the uncompressed file. We envisage our findings and framework serving as a benchmark in future development and analyses of lossy genomic data compressors.

The *Supplementary Data* can be found at http://web.stanford.edu/~iochoa/supplementEffectLossy.zip.

## I. Introduction

Recent advancements in Next Generation high-throughput Sequencing (NGS) have led to a drastic reduction in the cost of sequencing a genome (http://goo.gl/kKvmDl). This has generated an unprecedented amount of genomic data that must be stored, processed, and transmitted. To facilitate this effort, data compression techniques that allow for more efficient storage as well as fast exchange and dissemination of these data have been proposed in the literature.

Genome sequencing files, such as FASTQ and SAM/BAM, are mainly composed of nucleotide sequences (called *reads*) and quality scores that indicate the reliability of each of the nucleotides. According to SAM file specifications [1], the quality scores are stored using the *Phred score,* which corresponds to the number *Q* = − 10 log_10_ *P* (rounded to the closest integer), where *P* indicates the probability that the corresponding nucleotide in the read is in error. These scores are commonly represented using the ASCII alphabet [33 : 73] or [64 : 104], where the value corresponds to *Q* + 33 or *Q* + 64, respectively.

When losslessly compressed, quality scores comprise more than half of the compressed file [2]. In addition, it has been shown that the quality scores are inherently noisy [2], and downstream applications that use them do so in a heuristic manner. For these reasons, lossy compression of quality scores has been proposed to further reduce the storage requirements at the cost of introducing a distortion (i.e., the reconstructed quality scores may differ from the original ones).

Several lossy compressors of quality values have been proposed in the recent literature. These lossy compressors can be divided into two categories depending on whether or not they use biological information for the compression. While the majority of the proposed algorithms do not rely on such side information (see the survey on lossy compressors described in [3]), examples of compressors that do are in [4], [5].

Traditionally, lossy compressors have been analyzed in terms of their rate-distortion performance. Such analysis provides a yardstick for comparison of lossy compressors of quality scores that is oblivious to the multitude of downstream applications, which use the quality scores in different ways. However, the data compressed is used for biological inference. Researchers are thus more interested in understanding the effect that the distortion introduced in the quality scores has on the subsequent analysis performed on the data.

To date, there is not a standard practice on how this analysis should be performed. Proof of this is the variety of analyses presented in the literature when a new lossy compressor for quality scores is introduced (see [6], [7], [3], [5] and references therein). Moreover, it is not yet well understood how lossy compression of quality scores affects the downstream analysis performed on the data. This can be understood not only by the lack of a standard practice, but also by the variety of applications that exist and the different manner in which they use quality scores. In addition, the fact that lossy compressors can work at different rates and be optimized for several distortion metrics make the analysis more challenging. However, such an analysis is important if lossy compression is to become a viable mode for coping with the surging requirements of genomic data storage.

With this in mind, in this work we propose a methodology to analyze how lossy compression of quality scores affects the output of one of the most widely used downstream applications: variant calling, which compromises Single Nucleotide Polymorphism (SNP) and INDEL calling. Furthermore, we use the proposed methodology to compare the performance of the recently proposed lossy compressors for quality scores, which to our knowledge is the first in depth comparison available in the literature.

We focus on those lossy compressors that use only the quality scores for compression, as it would be too difficult to draw conclusions about the underlying source that generates the quality scores from analyzing algorithms like [5], where the lossy compression is done mainly using the information from the reads. Note also that in these cases it is not possible to specify a rate or a distortion to be minimized.

For the study, we compare the output of the variant caller when the quality scores of the original (uncompressed) data are replaced by the reconstructed ones. Specifically, for SNP calling we use the human dataset NA12878, which has been thoroughly characterized by GIAB [8], and for which a gold standard (consensus of SNPs) is available. To evaluate the effect of lossy compression on INDEL identification, we simulated genomes that contain biologically realistic SNPs and INDELs, creating a ground truth dataset. We then computationally generated sequencing reads for these genomes. In the following we assume these indels and the SNPs of the gold standard are the true ones and refer to them as the “ground truth”. This allows us to analyze which lossy compressor, distortion metric and rate produces the more accurate set of variants. We also show that in some cases applying lossy compression to the quality scores instead of lossless compression results in a set of variants that is more accurate. This suggests that lossy compression of quality scores can be beneficial not only for compression, but also to improve the inference performed on the data. The results presented in the manuscript also provide insight into the characteristics that a lossy compressor should have so that the reconstructed quality scores from which the set of variants is inferred do not differ much from those called based on the original quality scores.

We hope the methodology for variant calling analysis presented in this work will be of use in the future when introducing new lossy compressors. We leave the extensions of the investigations presented herein to other downstream applications for future work.

## II. Methodology for variant calling

In this section we describe the methods used to test the effect of lossy compressors on variant calling. The methodologies suggested for SNPs and indels differ, and thus we introduce each of them separately.

### A. SNP calling

Based on the most recent literature that compares different SNP calling pipelines ([9], [10], [11], [12], [13]) we have selected three pipelines for our study. Specifically, we propose the use of: i) the SNP calling pipeline suggested by the Broad Institute, which uses the Genome Analysis Toolkit (GATK) software package [14], [15], [16]; ii) the pipeline presented in the Hight Throughput Sequencing LIBrary (htslib.org), which uses the Samtools suit developed by The Wellcome Trust Sanger Institute [17]; and iii) the recently proposed variant caller named Platypus developed by Oxford University [18]. In the following we refer to these pipelines as **GATK**, **htslib.org**^1^ and **Platypus**, respectively.

In all pipelines we use BWA-mem [19] to align the FASTQ files to the reference (NCBI build 37, in our case), as stated in all best practices. For specific steps and the respective commands we refer the reader to the *Suplementary Data.*

Regarding the GATK pipeline, we note that the best practices recommends to further filter the variants found by the Haplotype caller by either applying the Variant Quality Score Recalibration (VQSR) or the Hard Filter. The VQSR filter is only recommended if the data set is big enough (more than 100K variants), since otherwise one of the steps of the VQSR, the Gaussian mixture model, may be inaccurate. Therefore, in our analysis we consider the use of both the VQSR and the Hard Filter after the Haplotype caller, both as specified in the best practices.

### B. INDEL detection

To evaluate the effect of lossy compression of base quality scores on INDEL calling, we employ popular INDEL detection pipelines: Pindel [20], Dindel [21], Unified Genotyper, Haplotype Caller [14], [15], [16] and Freebayes [22]. First, reads were aligned to the reference genome, NCBI build 37, with BWA [23]. We replaced the quality scores of the corresponding SAM/BAM file by those obtained after applying various lossy compressors, and then we performed the INDEL calling with each of the five tools. The *Supplementary Data* contains the detailed description of the commands necessary to run each pipeline. Note that several of these pipelines can be used to call both SNPs and INDELs, but the commands or parameters are different for each variant type.

### C. Datasets for SNP calling

A crucial part of the analysis is the ground truth, as it serves as the baseline for comparing the performance of the different lossy compressors against the lossless case. Thus, for the SNP calling analysis, we consider datasets from the *H. Sapiens* individual NA12878, for which two “ground truth” of SNPs have been released. In particular, we consider the datasets ERR174324 and ERR262997, which correspond to a 15 ×-coverage pair-end WGS dataset and a 30 ×-coverage pair-end WGS dataset, respectively. For each of them we extracted the chromosomes 11 and 20. Regarding the two “ground truths”, they are the one released by the Genome in a Bottle consortium (GIAB) [8], which has been adapted by the National Institute of Standardizations and Technology (NIST); and the ground truth released by Illumina as part of the Platinium Genomes project (http://www.illumina.com/platinumgenomes). Fig. 1 summarizes the differences between the two. As can be observed, most of the SNPs contained in the NIST ground truth are also included in Illumina’s ground truth, for both chromosomes. Note also that the number of SNPs on chromosome 20 is almost half of chromosome 11, for both “ground truths”^2^.

**Fig. 1.**
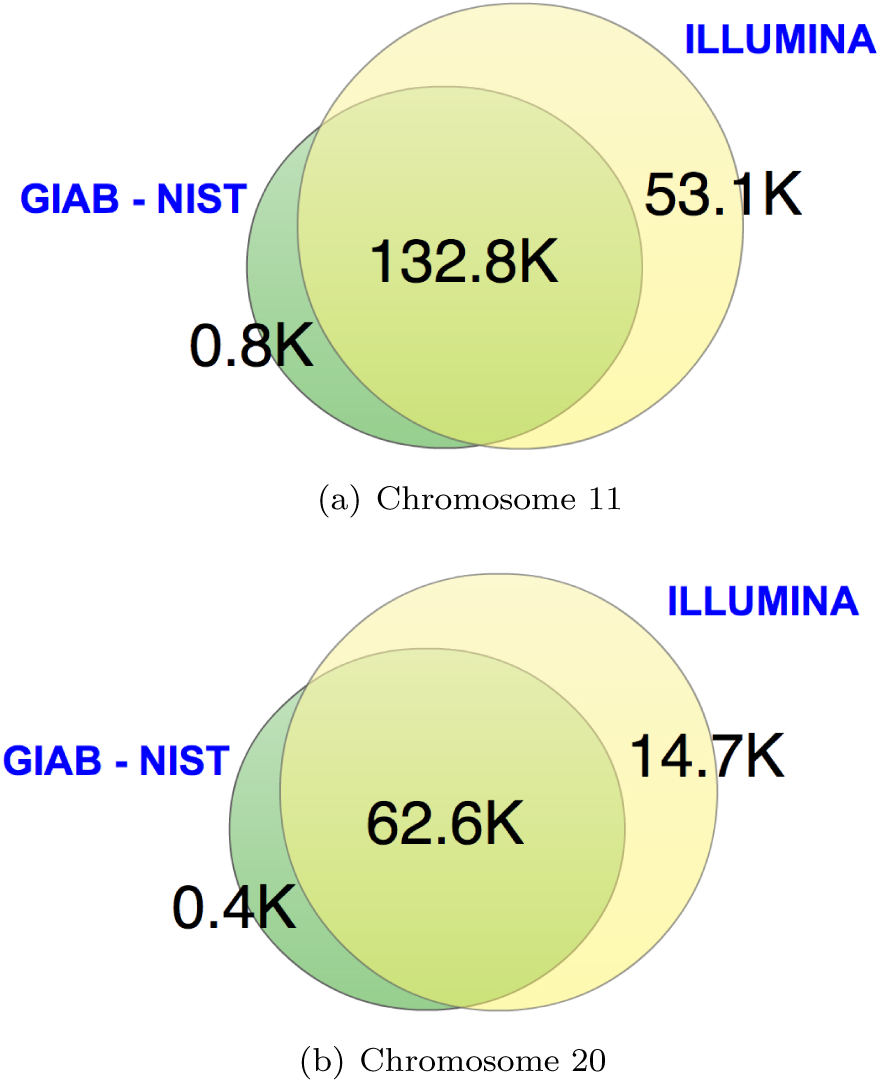
Difference between the GIAB NIST “ground truth” and the one from Illumina, for (a) chromosomes 11 and (b) 20.

### D. Datasets for INDEL detection

To evaluate the effect of lossy compression on INDEL detection, we simulated a chromosome containing 3000 heterozygous INDELs to use as a ground truth dataset. This approach gives us full knowledge and control of the exact location, zygosity, and alternative allele sequence of each INDEL, allowing us to robustly identify true positive and true negative variant calls. To best mimic true biological conditions, our simulated INDELs are drawn from biologically realistic size distributions, locations (coding vs non-coding), and insertion to deletion ratios seen in the Mills and 1000Genomes INDELs provided as part of the GATK bundle.

Using this simulated chromosome, we generated 100bp paired-end sequencing reads (using an Illumina-like error profile) with ART [24].

### E. Performance metrics

The output of each of the pipelines is a VCF file [25] which contains the set of the called variants. We first consider the case in which all the variants contained in the VCF file are positive calls, since the pipelines already follow their “best practices” to generate the corresponding VCF file. Since a ground truth exists for the datasets under consideration, we can compare these variants with those contained in the ground truth. Specifically, the comparison is made in terms of True Positives (T.P.), which refer to those variants contained both in the VCF file under consideration and the ground truth (a match in both position and genotyping must occur for the call to be declared T.P., while for INDELs the criteria were more lenient: any INDEL within 10bp of the true location was considered a T.P.); False Positives (F.P.), which refer to those variants contained in the VCF file under consideration but not in the ground truth; and False Negatives (F.N.), which correspond to those variants not contained in the VCF file under consideration but contained in the ground truth. The more T.P. (or equivalently the fewer F.N.) and the fewer F.P. the better. Ideally, we would like to apply a lossy compressor to the quality scores, such that the resulting file is smaller than that of the losslessly compressed, while obtaining a similar number of T.P. and F.P. We will show that not only is this possible, but that in some cases we can simultaneously obtain more T.P. and fewer F.P than with the original data.

To analyze the performance of the lossy compressors on the proposed pipelines, we employ the widely used metrics *sensitivity* and *precision,* which include in their calculation the true positives, false positives and false negatives, as described below:

- Sensitivity: measures the proportion of all the positives that are correctly called, computed as 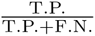.
- Precision: measures the proportion of called positives that are true, computed as 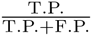.

Depending on the application, one may be inclined to boost the sensitivity at the cost of slightly reducing the precision, in order to be able to find as many T.P. as possible. On the other hand, there may be applications where it is more natural to optimize for precision than sensitivity. In an attempt to provide a measure that combines the previous two, we also use the f-score:

- F-score: the harmonic mean of the sensitivity and precision, computed as 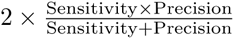.

In the discussion above we have considered that all the variants contained in a VCF file are positive calls. However, there is another common approach that consists of considering as positive calls only those that satisfy a given constraint, and as negative calls the remaining ones. In general, this constraint consists of having the value of one of the parameters associated with a variant above a certain threshold. This approach is used to construct the well known Receiver Operating Curves (ROC). In the case under consideration, the ROC curve shows the performance of the variant caller as a classification problem, that is, it shows how well the variant caller differentiates between true and false variants when filtered by a certain parameter. Specifically, it plots the False Positive Rate (F.P.R.) versus the True Positive Rate (T.P.R.) (also denoted as sensitivity) for all thresholding values. Given a ROC plot with several curves, a common method for comparing them is by calculating the area under the curve (AUC) of each of them, such that the larger the AUC the better.

The main drawback with this approach is the selection of the thresholding parameter. For instance, in SNP calling, when using the GATK pipeline, the *QD* (Quality by Depth) field is as valid a parameter as the *QUAL* field. Moreover, for different choices of the thresholding parameters, different performances of the AUC are obtained, as shown in the *Supplementary Data.* Given this, we believe that this approach is mainly suitable to analyze the VCF files that contain a clear thresholding parameter, like those VCF files obtained by the GATK pipeline after applying the VQSR filter, since in this case there is a clear parameter to be selected, namely the *VQSLOD*. Moreover, the considered pipelines already performed their own filtering as part of their “best practice” workflow.

## III. Lossy Compressors

To our knowledge, and based on the results presented in [3], RBlock, PBlock [7] and QVZ [3] are the algorithms that perform better among the existing lossy compressors that solely use the quality scores to compress. Therefore, those are the algorithms that we consider for our study. In addition, we consider Illumina’s proposed binning (http://goo.gl/d5TYDk), which is implemented both by CRAM and DSRC2. In the Results section we refer to the performance of DSRC2 [26] rather than CRAM [27], as it generates a compressed file of smaller size (see [3]). Next, we describe the aforementioned lossy compressors in more detail.

**P/R-Block:** The *PBlock* and *RBlock* algorithms were introduced in [7]. Both algorithms represent quality scores by separating them into blocks of variable size, such that all the quality scores contained in a block can be replaced by the same representative value without violating a given distortion constraint. The algorithms then store for each block its length and the representative value, which are losslessly compressed. What differs between the algorithms is the distortion constraint, that we specify next.

Given a block of quality scores, *Q*_max_ and *Q*_min_ denote the largest and smallest quality scores within the block, respectively. In PBlock, the quality scores contained in a block should satisfy *Q*_max_ − *Q*_min_ ≤ 2*p,* where *p* is a user specified parameter. On the other hand, in RBlock the quality scores contained in a block should satisfy 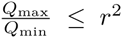, where *r* is a user specified parameter.

The main difference between the two algorithms is related to the maximum absolute distance allowed between a quality score and its representative (the new quality score). Whereas in PBlock this distance is constant for every quality score, in RBlock a low quality score will in general be closer to its representative than a high quality score. That is, the algorithm is more precise in representing low quality scores than high ones. Finally, note that in both algorithms the maximum absolute distance between a quality score and its representative is controlled by the user.

**QVZ - Quality Values Zip:** QVZ was introduced in [3], and it allows the user to choose the rate and the distortion to be minimized (the built in distortions are Mean Square Error (MSE), Lorentzian and L1). QVZ assumes a Markov model of order 1 for compression, and it computes the statistics at each position empirically from the data. In brief, given those statistics, the distortion to be minimized and the rate, the algorithm makes use of the Lloyd-Max algorithm [28] to compute the best quantizers. Note that at each position there are as many quantizers as different values in the previous position (due to the Markov assumption). A quantizer is composed of decision regions and the representatives of each region. Once all the quantizers are computed, QVZ assigns each quality score to the corresponding decision region, which is then losslessly compressed by an adaptive arithmetic encoder. In order to improve the rate-distortion performance, QVZ has the option of clustering the data prior to compression.

**Illumina’s Binning:** Illumina’s proposed binning reduces the alphabet size by applying an 8 level mapping. The specific mapping is summarized in Table I. As can be inferred from the table, the applied mapping is more precise in representing high quality scores than low ones (based on the size of the bins). Also, note that the maximum distance between an original quality score and the new one is always upper bounded by 5.

**TABLE I.** Illumina’s proposed 8 level mapping.

| Quality Score Bins | New Quality Score |
| --- | --- |
| N (no call) | N (no call) |
| 2-9 | 6 |
| 10-19 | 15 |
| 20-24 | 22 |
| 25-29 | 27 |
| 30-34 | 33 |
| 35-39 | 37 |
| $\geq 40$ | 40 |

### A. Comparison of lossy compressors

There are some important differences between the lossy compressors introduced above. For example, the compression scheme of Illumina’s proposed binning does not depend on the statistics of the quality scores, whereas QVZ and P/R-Block do. Also, in both Illumina’s proposed binning and P/R-Block the maximum absolute distance between a quality score and its reconstructed one (after decompression) can be controlled by the user, whereas in QVZ this is not the case. The reason is that QVZ designs the quantizers to minimize a given average distortion based on a rate constraint, and thus even though on average the distortion is small, some specific quality scores may have a reconstructed quality score that is far from the true one. Also, note that whereas Illumina’s proposed binning applies more precision to high quality scores, R-Block does the opposite, and P-Block does it equally among all the quality scores. Finally, in Illumina’s proposed binning and P/R-Block the user cannot estimate the size of the compressed file in advance, whereas this is possible in QVZ.

## IV. Results

We analyze the output of the variant caller (i.e., the VCF file) for each of the introduced pipelines when the quality scores are replaced by those generated by a lossy compressor. We focus on the following lossy compressors: QVZ, PBlock, RBlock and DSRC2. Recall that in QVZ we can choose the distortion, the rate and the number of clusters, and in PBlock and RBlock the parameters *p* and r, respectively. DSRC2 uniquely performs Illumina’s proposed binning. Thus, except for DSRC2, we run each of the algorithms several times with different parameters, generating different quality scores for each run. Specifically, we used QVZ with 1 and 3 clusters, rates ranging from 0 to 1, and the three built-in distortions MSE, L1 and Lorentzian (we refer to them as M, A and L, respectively). For PBlock we considered values of *p* ranging from 1 to 32, and for RBlock values of *r* ranging from 3 to 30.

Due to space constraints, here we show the results for QVZ with MSE distortion and three clusters, denoted as *QVZ-Mc3,* RBlock and the Illumina proposed binning. We selected these as they are good representatives of the overall results. We refer the reader to the *Supplementary Data* for the results with all the aforementioned parameters.

### A. SNP calling

We show that it is possible to perform lossy compression on the quality values, while simultaneously increasing the number of true positives and decreasing the number of false positives with respect to the original dataset, when using a variant calling pipeline that follows its “best practice” to generate the corresponding VCF file. Fig. 2 shows the behavior obtained by the different lossy compressors in the GATK pipeline with hard filtering with the ERR262996 dataset (30× coverage), chromosome 11, when the ground truth is given by that of Illumina (similar plots for the remaining cases are shown in the *Supplementary Data*). For ease of visualization, we normalize the number of T.P. and F.P. with those generated with the uncompressed file, such that the number of T.P. and F.P. of the uncompressed VCF after normalization become 0. We then join the points that belong to a given algorithm, sorted by size, such that the point closer to (0,0) corresponds to the largest size. As is evident from the figure, several points have a positive number of T.P. and a negative number of F.P., which corresponds to an improvement over the uncompressed one. Moreover, in this case storage of the quality scores is reduced from 1.3 GB (when lossless compressed with QVZ) to 0.25 GB, which corresponds to more than 80% reduction. As shown in the *Supplementary Data* and in the results that follow, this phenomenon is not idiosyncratic to this particular dataset and pipeline, but in fact widely observed in the other datasets and pipelines as well.

**Fig. 2.**
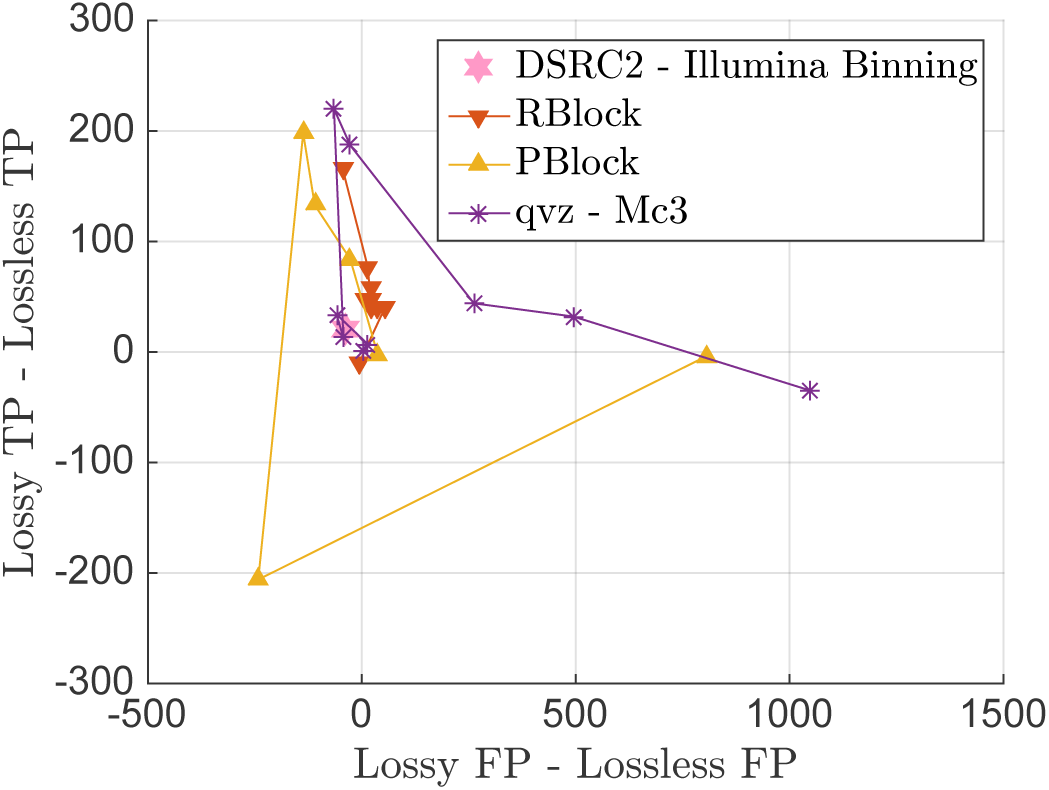
T.P. vs F.P., with respect to illumina’s ground truth, for the GATK pipeline and 30× coverage dataset ERR262996, chromosome 11. Different points within a given lossy compressor correspond to the use of different parameters (and thus compression rate). For ease of visualization, we added a line that joins the different points from smaller to higher rate, where the point furthest from (0,0) is the one with lowest rate.

As mentioned in the methodology, we start by showing the value of the sensitivity, precision and f-score achieved by each of the algorithms for each of the pipelines, dataset and ground truth, together with the compression ratio (e.g., a reduction from 100MB to 80MB corresponds to a 20% compression ratio). The latter is computed with respect to the lossless compressed quality scores, computed using QVZ in lossless mode (which, as shown in [3], performs similarly to the state-of-the-art lossless compressors for quality scores). We choose to show the results using tables as they help visualize which lossy compressors and/or parameters work better for a specific setting. We color in red (will appear as a shaded cell) the values of the sensitivity, precision and f-score that improve upon the uncompressed. We also generated a table for each pipeline and/or ground truth that contains the average behavior of each of the algorithms with the different data sets. Due to the large amount of tables and their size, we provide excel files (.xlsx) as *Supplementary Data* that contain all the generated tables.

Table II, III and IV show the results for algorithms RBlock, QVZ-Mc3 (MSE distortion criteria and 3 clusters) and Illumina binning-DSRC2 for the GATK with hard filtering, htslib.org and Platypus pipelines, respectively, when using the NIST ground truth. The two columns refer to the average results of Chromosomes 11 and 20 of the ERR262996 and ERR174310 datasets, respectively. We refer to the *Supplementary Data (.xlsx)* for the results of QVZ using other distortions and rates, and for PBlock, as well as for the results of individual chromosomes.

**TABLE II.** Sensitivity, Precision, F-score and Compression ratio for the 30× and 15× Coverage datasets for the GATK pipeline, using the NIST ground truth.

| GATK | ERR262996 (30×): Chr11,Chr20 |  |  |  | ERR174310 (15×): Chr11,Chr20 |  |  |  |
| --- | --- | --- | --- | --- | --- | --- | --- | --- |
|  | Sensitivity | Precision | F-Score | Compression | Sensitivity | Precision | F-Score | Compression |
| Lossless | 0.9829 | 0.7422 | 0.8456 | 0 | 0.9699 | 0.7332 | 0.8351 | 0 |
| Rblock |  |  |  |  |  |  |  |  |
| 3 | 0.9828 | 0.7424 | 0.8457 | -16.7 | 0.97 | 0.7331 | 0.835 | -16.9 |
| 8 | 0.983 | 0.7421 | 0.8456 | 31 | 0.9708 | 0.733 | 0.8353 | 31.6 |
| 10 | 0.983 | 0.7422 | 0.8457 | 42 | 0.9708 | 0.733 | 0.8353 | 43 |
| 20 | 0.9831 | 0.7422 | 0.8457 | 67.4 | 0.971 | 0.7328 | 0.8352 | 70.7 |
| 30 | 0.9833 | 0.7422 | 0.8458 | 76.4 | 0.9709 | 0.7326 | 0.8351 | 78.9 |
| QVZ-Mc3 |  |  |  |  |  |  |  |  |
| 0.9 | 0.9829 | 0.7421 | 0.8456 | 10.2 | 0.9699 | 0.7332 | 0.8351 | 10 |
| 0.8 | 0.983 | 0.7425 | 0.8459 | 14.7 | 0.9699 | 0.7333 | 0.8351 | 19 |
| 0.6 | 0.9828 | 0.7424 | 0.8457 | 37.6 | 0.97 | 0.7326 | 0.8347 | 38.3 |
| 0.4 | 0.9833 | 0.7422 | 0.8458 | 57.5 | 0.9691 | 0.7321 | 0.8341 | 58.6 |
| 0.2 | 0.9831 | 0.7418 | 0.8454 | 77.7 | 0.9679 | 0.7309 | 0.8328 | 77.9 |
| Illumina-DSRC2 | 0.9827 | 0.7424 | 0.8457 | 54.78 | 0.9694 | 0.7345 | 0.8357 | 55.66 |

**TABLE III.**
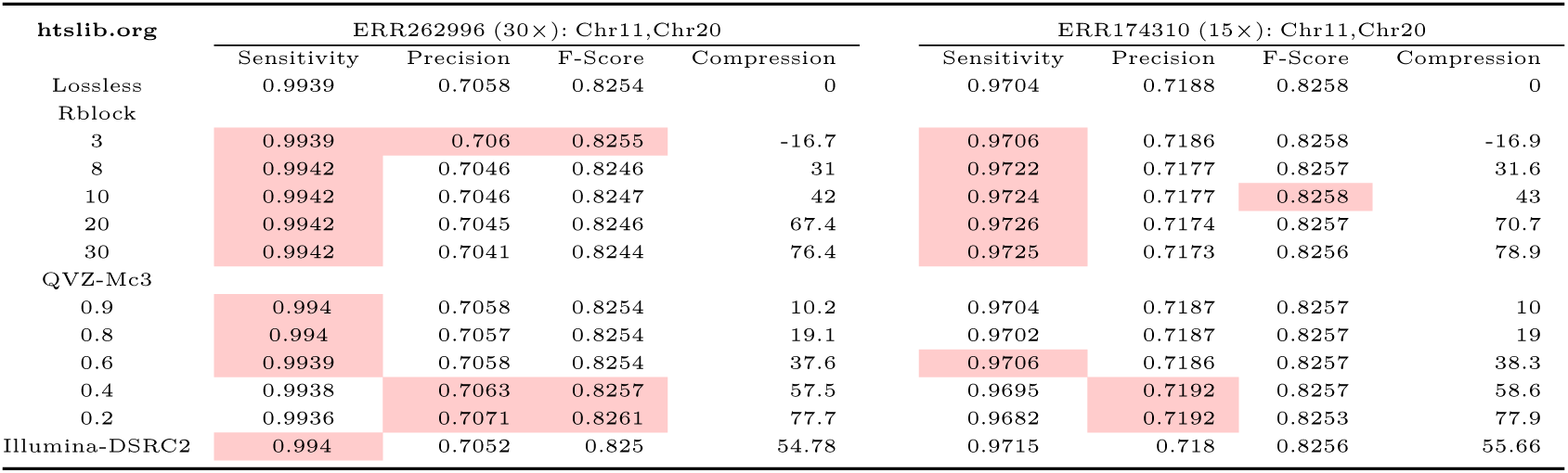
Sensitivity, Precision, F-score and Compression ratio for the 30× and 15× Coverage datasets for the htslib.org pipeline, using the NIST ground truth.

| htslib.org | ERR262996 (30×): Chr11,Chr20 |  |  |  | ERR174310 (15×): Chr11,Chr20 |  |  |  |
| --- | --- | --- | --- | --- | --- | --- | --- | --- |
|  | Sensitivity | Precision | F-Score | Compression | Sensitivity | Precision | F-Score | Compression |
| Lossless | 0.9939 | 0.7058 | 0.8254 | 0 | 0.9704 | 0.7188 | 0.8258 | 0 |
| Rblock |  |  |  |  |  |  |  |  |
| 3 | 0.9939 | 0.706 | 0.8255 | -16.7 | 0.9706 | 0.7186 | 0.8258 | -16.9 |
| 8 | 0.9942 | 0.7046 | 0.8246 | 31 | 0.9722 | 0.7177 | 0.8257 | 31.6 |
| 10 | 0.9942 | 0.7046 | 0.8247 | 42 | 0.9724 | 0.7177 | 0.8258 | 43 |
| 20 | 0.9942 | 0.7045 | 0.8246 | 67.4 | 0.9726 | 0.7174 | 0.8257 | 70.7 |
| 30 | 0.9942 | 0.7041 | 0.8244 | 76.4 | 0.9725 | 0.7173 | 0.8256 | 78.9 |
| QVZ-Mc3 |  |  |  |  |  |  |  |  |
| 0.9 | 0.994 | 0.7058 | 0.8254 | 10.2 | 0.9704 | 0.7187 | 0.8257 | 10 |
| 0.8 | 0.994 | 0.7057 | 0.8254 | 19.1 | 0.9702 | 0.7187 | 0.8257 | 19 |
| 0.6 | 0.9939 | 0.7058 | 0.8254 | 37.6 | 0.9706 | 0.7186 | 0.8257 | 38.3 |
| 0.4 | 0.9938 | 0.7063 | 0.8257 | 57.5 | 0.9695 | 0.7192 | 0.8257 | 58.6 |
| 0.2 | 0.9936 | 0.7071 | 0.8261 | 77.7 | 0.9682 | 0.7192 | 0.8253 | 77.9 |
| Illumina-DSRC2 | 0.994 | 0.7052 | 0.825 | 54.78 | 0.9715 | 0.718 | 0.8256 | 55.66 |

**TABLE IV.** Sensitivity, Precision, F-score and Compression ratio for the 30× and 15× Coverage datasets for the Platypus pipeline, using the NIST ground truth.

| Platypus | ERR262996 (30×): Chr11,Chr20 |  |  |  | ERR174310 (15×): Chr11,Chr20 |  |  |  |
| --- | --- | --- | --- | --- | --- | --- | --- | --- |
|  | Sensitivity | Precision | F-Score | Compression | Sensitivity | Precision | F-Score | Compression |
| Lossless | 0.9368 | 0.7261 | 0.818 | 0 | 0.921 | 0.7426 | 0.822 | 0 |
| Rblock |  |  |  |  |  |  |  |  |
| 3 | 0.9368 | 0.7263 | 0.8181 | -16.7 | 0.9206 | 0.7427 | 0.822 | -16.871 |
| 8 | 0.9367 | 0.7268 | 0.8184 | 31 | 0.9202 | 0.7431 | 0.822 | 31.5852 |
| 10 | 0.9367 | 0.7268 | 0.8184 | 42 | 0.9201 | 0.7431 | 0.822 | 42.9952 |
| 20 | 0.9365 | 0.7276 | 0.8188 | 67.4 | 0.9193 | 0.7434 | 0.8219 | 70.685 |
| 30 | 0.9363 | 0.7277 | 0.8188 | 76.4 | 0.9184 | 0.7436 | 0.8216 | 78.9077 |
| QVZ-Mc3 |  |  |  |  |  |  |  |  |
| 0.9 | 0.9368 | 0.7262 | 0.8181 | 10.2 | 0.9205 | 0.7427 | 0.8219 | 10.0149 |
| 0.8 | 0.9368 | 0.7262 | 0.8180 | 19.1 | 0.9206 | 0.7427 | 0.8219 | 19.0204 |
| 0.6 | 0.9368 | 0.7265 | 0.8182 | 37.6 | 0.9206 | 0.7428 | 0.8220 | 38.2550 |
| 0.4 | 0.9369 | 0.725 | 0.8173 | 57.5 | 0.9206 | 0.7406 | 0.8206 | 58.5607 |
| 0.2 | 0.9368 | 0.7219 | 0.8153 | 77.7 | 0.9206 | 0.734 | 0.8166 | 77.9354 |
| Illumina-DSRC2 | 0.9368 | 0.7261 | 0.8179 | 54.78 | 0.9208 | 0.7425 | 0.8219 | 55.6 |

It is worth noting that with the GATK pipeline, several points improve simultaneously the sensitivity, precision and f-score when compared to the uncompressed (original) quality scores. For example, in the 30× coverage dataset, RBlock improves the performance while reducing the size by more than 76% (PBlock manages to boost the compression to more than 80%, see *Supplementary Data (.xlsx)).* In the 15× coverage dataset QVZ improves upon the uncompressed and reduces its size by 20%. With the htslib.org pipeline, it is interesting to see that most of the points improve the sensitivity parameter, meaning that they are able to find more T.P. than with the uncompressed quality scores. As shown in the *Supplementary Data (.xlsx),* QVZ with L1 and Lorentzian distortion achieves better performance than the uncompressed (in all three metrics) while reducing the quality scores by more than 10%, for the 30× coverage dataset. Finally, with the Platypus pipeline, the parameters that improve in general are the precision and the f-score, which indicates that a bigger percentage of the calls are T.P. rather than F. P. Some points also improve upon the uncompressed, like QVZ with L1 distortion, which achieves almost 38% compression (in the 30× coverage dataset).

Similar tables when the ground truth is provided by Illumina are contained in the *Supplementary Data (.xlsx).* In that case, with the GATK pipeline, R/P-Block improves mainly the sensitivity and f-score, with PBlock improving the precision as well in the 30× coverage dataset. QVZ seems to perform better in this case, improving upon the uncompressed for several rates. It also achieves a performance better than that of Illumina’s proposed binning for a similar compression rate. With the htslib.org pipeline R/P-Block improve mainly the sensitivity, while QVZ improves the precision and the f-score (in the 30× coverage dataset). The performance on Platypus is similar to the one obtained when the NIST ground truth is used instead.

In summary, the performance of QVZ with 3 clusters is in general better than with 1 cluster, especially for small rates. In terms of the distortion metric that QVZ aims to minimize, MSE works significantly better for small rates (in most of the cases), whereas for higher rates the three analyzed distortions offer a similar performance. Thus the compression rate seems much more significant to the variability in the performance than the choice of distortion criterion. RBlock offers in general better performance than PBlock for similar compression rates. Finally, in most of the analyzed cases, Illumina’s binning is outperformed by at least one other lossy compressor, while offering a similar compression rate. Overall, for high compression ratios (30%-70%), RBlock seems to perform the best, whereas QVZ is preferred for lower compression rates (>70%).

To get more insight into the possible benefits of using lossy compression, we show the distribution of the f-score difference between the lossy and lossless case for different lossy compressors and rates (thus a positive number indicates an improvement over the lossless case). The distribution is computed by averaging over all simulations (24 values in total). Figure 3 shows the box-plot and the mean value of the f-score difference for six different compression rates. Since QVZ performs better for low compression rates, we show the results for QVZ-Mc3 with parameters 0.9, 0.8 and 0.6 (left-most side of the figure). Analogously, for high compression ratios we show the results of RBlock with parameters 30, 20, and 10 (rightmost side of the figure).

**Fig. 3.**
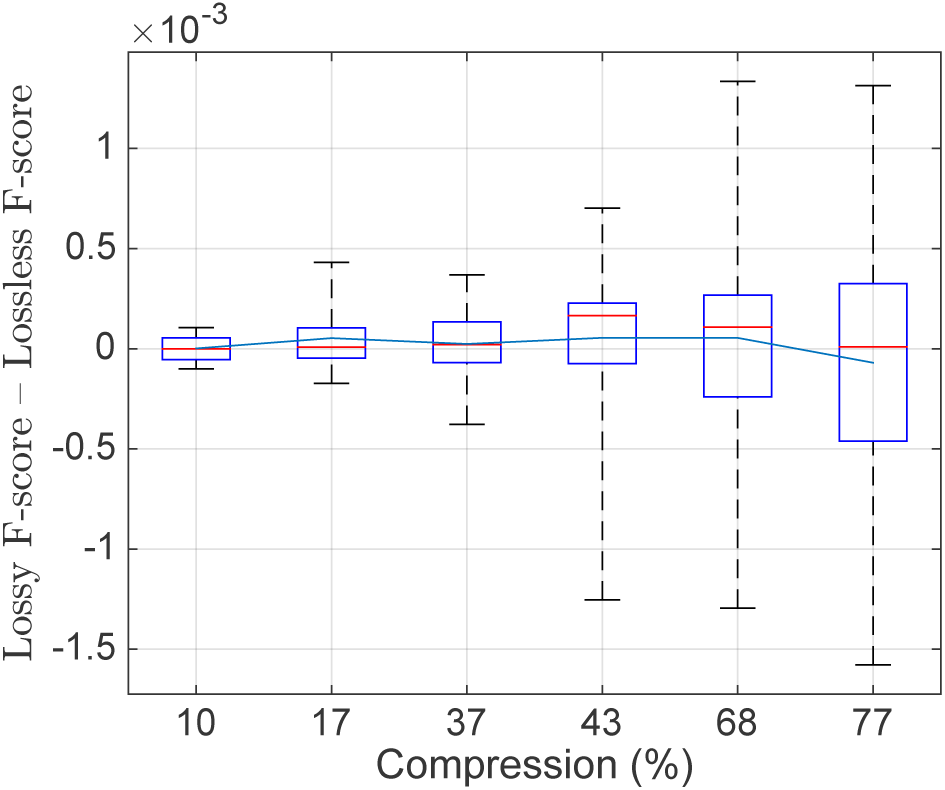
Box plot of f-score differences between the lossless case and six lossy compression algorithms. The x-axis shows the compression rate achieved by the algorithm. The three left-most boxes correspond to QVZ-Mc3 with parameters 0.9, 0.8 and 0.6; while the three rightmost boxes correspond to RBlock with parameters 30, 20 and 10. The blue line indicates the mean value, and the red one the median.

Remarkably, for all the rates the median is positive, which indicates that in at least 50% of the cases lossy compression improved upon the uncompressed quality scores. Moreover, the mean is also positive, except for the point with highest compression. This suggests that lossy compression may be used to reduce the size of the quality scores without compromising the performance on the SNP calling. These findings are in line with the results provided above, where we showed that in several cases the sensitivity, precision and f-score could be improved simultaneously while significantly reducing the quality score storage requirements.

In the previously analyzed cases we have assumed that all the SNPs contained in the VCF file are positive calls, since the pipelines already follow their “best practice” to generate the corresponding VCF file. As discussed in the Methodology, another possibility is to select a parameter and consider positive calls only those whose parameter is above a certain threshold. Varying the threshold results in the ROC curve. We believe this approach is of interest to analyze the VCF files generated by the GATK pipeline followed by the VQSR filter, with thresholding parameter given the VQSLOD field, and thus we present the results for this case. For completeness, we also generated the ROC curves of the remaining cases (see *Supplementary Data*). Fig. 4 shows the ROC curve of chromosome 11 of the 30× coverage dataset (ERR262996), with the NIST ground truth. The results correspond to those obtained when the quality scores are the original ones (lossless), and the ones generated by QVZ-Mc3 (MSE distortion and 3 clusters), PBlock with parameter 8, RBlock with parameter 25 and the Illumina binning (as the results of applying the DSRC2 algorithm). As shown in the figure, each of the algorithms outperform the rest in at least one point of the curve. This is not the case for the Illumina Binning, as it is outperformed by at least one other algorithm in all points. Moreover, the AUC of all the lossy compressors except that of the Illumina Binning outperform that of the lossless case.

**Fig. 4.**
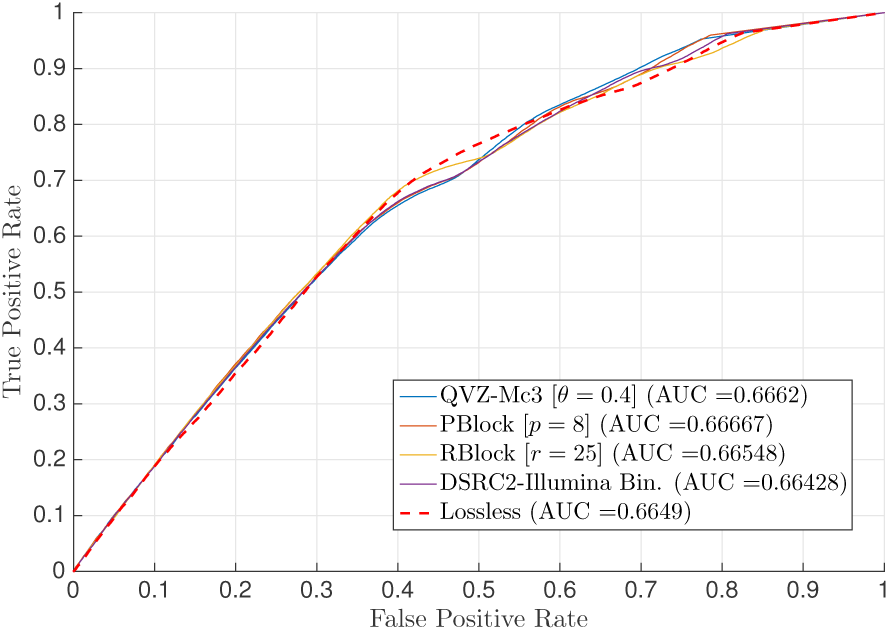
ROC curve of chromosome 11 (ERR262996) with the NIST ground truth and the GATK pipeline with the VQSR filter. The ROC curve was generated with respect to the VQSLOD field. The results are for the original quality scores (uncompressed), and those generated by QVZ-Mc3 (MSE distortion and 3 clusters), PBlock (*p* = 8) and RBlock (*r* = 25).

### B. INDEL calling

Similar to SNP calling, we show that lossy compression can lead to files of smaller size while obtaining INDEL detection accuracy that is similar to that obtained with the uncompressed data.

We observe that Pindel does not use the quality scores to perform the INDEL calling, and thus the results obtained with the data previously lossily compressed and the uncompressed are exactly the same (see *Supplementary Data (indels.xls)*). Therefore, in the following discussion we focus on the remaining pipelines. Table V displays results for Dindel and Unified Genotyper pipelines (see Supplementary Data for results for other lossy compressors). Shadowed in red are the points that outperform the results obtained with the uncompressed data. We observe that Rblock has higher sensitivity, precision, and F-score for identifying INDELs than the uncompressed data. Notice that the file size in this case increases with respect to the uncompressed. We believe the file size could be reduced by improving the entropy encoder of the Rblock algorithm. With QVZ it is possible to reduce the size by 10% while still improving with respect to the uncompressed. Finally, Illumina’s proposed binning maintains the precision, but obtains smaller values for both the sensitivity and the f-score. Unified Genotyper gives similar results to those from Dindel. The main differences are that in this case Pblock outperforms the uncompressed while obtaining a reduction in size of 29%, and Illumina is not able to improve with respect to the uncompressed (see *Supplementary Data*).

**TABLE V.**
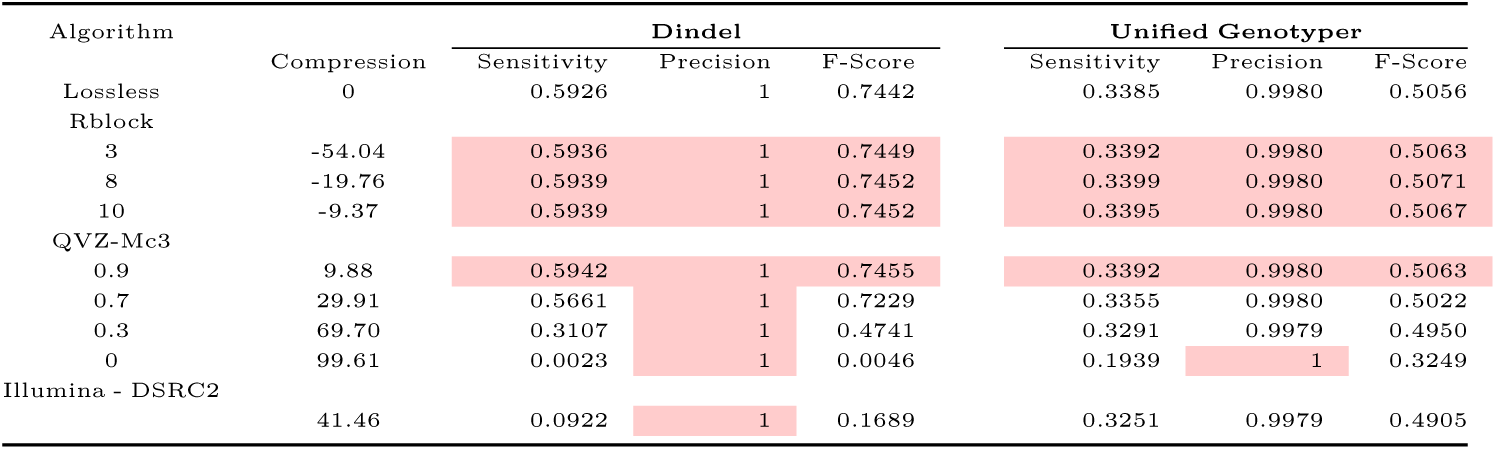
Sensitivity, Precision and F-score for the first dataset for the Dindel and Unified Genotyper pipelines.

| Algorithm | Compression | Dindel |  |  | Unified Genotyper |  |  |
| --- | --- | --- | --- | --- | --- | --- | --- |
|  |  | Sensitivity | Precision | F-Score | Sensitivity | Precision | F-Score |
| Lossless | 0 | 0.5926 | 1 | 0.7442 | 0.3385 | 0.9980 | 0.5056 |
| Rblock |  |  |  |  |  |  |  |
| 3 | -54.04 | 0.5936 | 1 | 0.7449 | 0.3392 | 0.9980 | 0.5063 |
| 8 | -19.76 | 0.5939 | 1 | 0.7452 | 0.3399 | 0.9980 | 0.5071 |
| 10 | -9.37 | 0.5939 | 1 | 0.7452 | 0.3395 | 0.9980 | 0.5067 |
| QVZ-Mc3 |  |  |  |  |  |  |  |
| 0.9 | 9.88 | 0.5942 | 1 | 0.7455 | 0.3392 | 0.9980 | 0.5063 |
| 0.7 | 29.91 | 0.5661 | 1 | 0.7229 | 0.3355 | 0.9980 | 0.5022 |
| 0.3 | 69.70 | 0.3107 | 1 | 0.4741 | 0.3291 | 0.9979 | 0.4950 |
| 0 | 99.61 | 0.0023 | 1 | 0.0046 | 0.1939 | 1 | 0.3249 |
| Illumina - DSRC2 | 41.46 | 0.0922 | 1 | 0.1689 | 0.3251 | 0.9979 | 0.4905 |

Table VI shows the results for the remaining pipelines, that is, Haplotype Caller and Freebayes. With the Haplotype Caller more true positives are found, but also more false positives, so only sensitivity improves. For Freebayes the results are minimally impacted by varying the compression approach, though, the results are improved with respect to the uncompressed. The only point for which this is not true is Illumina proposed binning, which calls only one variant that is a false positive.

**TABLE VI.**
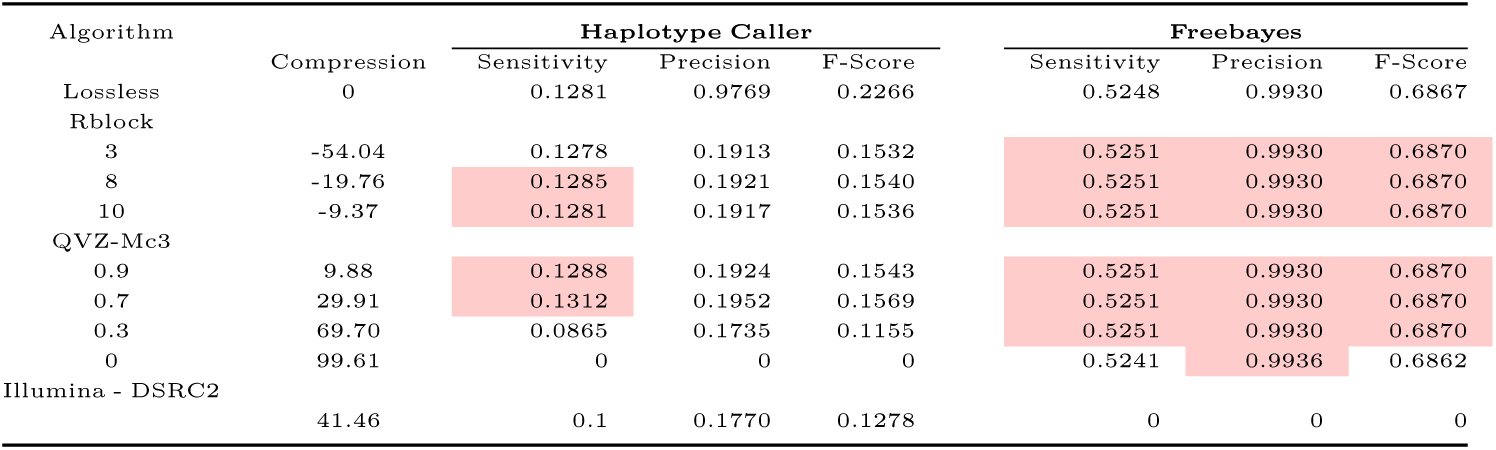
Sensitivity, Precision and F-score for the first dataset for the Haplotype Caller and Freebayes pipelines.

## V. Discussion

We have shown that lossy compressors can reduce file size at a minimal cost - or even benefit - to sensitivity and precision in SNP and INDEL detection.

Based on the results shown in the previous section, we conclude that in many cases lossy compression can significantly reduce the genomic file sizes (with respect to the losslessly compressed) without compromising the performance on the variant calling. Moreover, in several cases we have observed that lossy compression actually leads to results that improve over the uncompressed ones, i.e., they generate more true positives and fewer false positives than with the original quality scores, when compared to the corresponding ground truth. This behavior is consistent with observations from the recent literature (see for example [3], [5]).

We have analyzed several lossy compressors introduced recently in the literature. The main difference among them relates to the way they use the statistics of the quality scores for compression. For example, Illumina’s proposed binning is a fixed mapping that does not use the underlying properties of the quality scores. In contrast, algorithms like QVZ are fully based on the statistics of the quality scores to design the corresponding quantizers for each case. As manifested in the results, Illumina’s binning is generally outperformed by the other lossy compressors. This suggests that using the statistics of the quality scores for compression is beneficial, and that not all datasets should be treated in the same way.

Our findings put together with the fact that, when losslessly compressed, quality scores comprise more than 50% of the compressed file [2], seem to indicate that lossy compression of quality scores could become an acceptable practice in the future for boosting compression performance or when operating in bandwidth constrained environments. The main challenge in such a mode may be to decide which lossy compressor and/or rate to use in each case. Part of this is due to the fact that the results presented so far are experimental, and we have yet to develop theory that will guide the construction or choice of compressors geared toward improved inference. One reason is that the statistics of the noise inherent in the quality scores have yet to be understood and thus it is not possible to design lossy compressors tailored to them. Moreover, the results that show that lossy compression can lead to inference that improves upon the uncompressed suggest that the data could be denoised. In that regard, an understanding of the statistical characteristics of the noise would enable the design of denoisers/compressors that remove part of the noise as they compress (see for example [29]), thus improving the subsequent analysis performed on it.

Evidently, for lossy compression of quality scores to become a standard practice, further research is called for. It should include improved modeling of the statistics of the noise, construction of lossy compressors and denoisers tuned to such models, and more experimentation on real data with additional downstream applications. Further, the phenomenon observed here where lossy compression of the quality scores can actually boost the performance of the downstream applications is highlighting not only the potential in lossy compression of quality scores, but also the need for revisiting the design of the downstream applications to make more principled use of the quality scores (with and without compression).

## VI. Conclusion

Recently there has been a growing interest in lossy compression of quality scores as a way to reduce raw genomic data storage costs. However, the genomic data under consideration is used for biological inference, and thus it is important to first understand the effect that lossy compression has on the subsequent analysis performed on it. To date, there is no clear methodology to do so, as can be inferred from the variety of analyses performed in the literature when new lossy compressors are introduced. To alleviate this issue, in this paper we have described a methodology to analyze the effect that lossy compression of quality scores has on variant calling, one of the most widely used downstream applications in practice. We hope the described methodology will be of use in the future when analyzing new lossy compressors and/or new datasets.

Specifically, the proposed methodology considers the use of different pipelines for SNP calling and indel calling, and datasets for which true variants exist (“ground truth”). We have used this methodology to analyze the behavior of the state-of-the-art lossy compressors, which to our knowledge constitutes the most complete analysis to date. The results demonstrate the potential of lossy compression as a means to reduce the storage requirements while obtaining performance close to that based on the original data. Moreover, in many cases we have shown that it is possible to improve upon the original data, corroborating the belief that the quality scores are noisy and thus they can be denoised (in our case via compression).

Our findings and the growing need for reducing the storage requirements suggest that lossy compression may be a viable mode for storing quality scores. However, further research should be performed to better understand the statistical properties of the quality scores, as well as the noise underlying their generation, to enable the principled design of lossy compressors and/or denoisers tailored to them. Moreover, methodologies for the analysis on other important downstream applications should be developed.

## VII. Acknowledgement

*a) Funding:* This work is partially funded by a Stanford Graduate Fellowship Program in Science and Engineering, a fellowship from the Basque Government, a grant from the Center for Science of Information (CSol), the 2014-07364-01 NIH grant, the 1 U01 CA198943-01 NIH grant, the 1157849-1-QAZCC NSF grant, the National Library of Medicine Training Grant T15 LM7033 and an NSF graduate research fellowship.

*b) Conflict of Interest:* None declared.

## Footnotes

1 More commonly referred to as samtools.

2 As is clear from the discussion in this subsection, the term *ground truth* should be taken with a grain of salt and as such should appear in quotation marks throughout. We omit these marks henceforth for simplicity.

